# Role of acetoacetyl-CoA synthetase in glucose uptake by HepG2 cells

**DOI:** 10.1101/2023.02.04.527056

**Authors:** Shinya Hasegawa, Masahiko Imai, Noriko Takahashi

## Abstract

Accelerated glucose metabolism is a common feature of cancer cells and represents a potential therapeutic target. Recent studies have revealed that lipid metabolism is related to glucose me-tabolism, especially glucose uptake. Acetoacetyl-CoA synthetase (AACS) converts acetoacetate, a ketone body, to acetoacetyl-CoA, which is incorporated into cholesterol and fatty acids. The AACS gene is highly expressed in HepG2 cells but not in the human normal liver. These results suggest a correlation between AACS and glucose metabolism. Therefore, we examined the relationship between AACS and glucose uptake in hepatocellular carcinoma (HepG2) cells. The expression of AACS was significantly upregulated when HepG2 cells were exposed to high concentrations of glucose. Lentiviruses, which code for shRNA against AACS (shAACS), were used to determine whether AACS knockdown affects the glucose uptake in HepG2 cells. AACS knockdown signifi-cantly reduced glucose uptake and increased the concentration of ketone bodies in the media, and treatment with β-hydroxybutyrate, a ketone body, attenuated glucose uptake. Moreover, the knockdown of AACS, i.e., the increased concentration of the intracellular ketone body, slightly attenuated cell proliferation and significantly increased the sensitivity to the anticancer agent sorafenib. These results suggest that AACS plays an important role in glucose uptake and cell proliferation and that ketone bodies may compete with glucose as an important energy source for cancer cells.

## Introduction

Cancer is one of the leading causes of mortality worldwide. Its treatment is complicated, in part, because of the genetic diversity of cancer cells. A common feature of cancer cells is the reprograming of cellular metabolism that is a consequence of oncogenic mutation [1]. One well-known metabolic change is the Warburg effect in which cancer cells generate ATP via glycolysis rather than oxidative phosphorylation, even in the presence of oxygen. This accelerated glucose metabolism is used as a diagnostic marker. Furthermore, it has been proposed that key lipid metabolism genes may be used as prognostic biomarkers in several types of cancer associated with tumor recurrence and/or survival [2]. Thus, cancer cells have a metabolic profile that may be exploited to identify potential therapeutic targets and diagnostic markers.

Ketone bodies are considered an alternative energy source not only in normal tissues but also in cancer cells [3]. Several recent studies have shown that ketone body utilization drives tumor growth and metastasis in cancer cells [4]. Ketone bodies are utilized for energy production via succinyl-CoA:3-ketoacid-CoA transferase (EC 2.8.3.5) in the mitochondria [5,6]. We previously reported that acetoacetyl-CoA synthetase (AACS, acetoacetate-CoA ligase, EC 6.2.1.16), which is the other ketone body-utilizing enzyme, converts acetoacetate, a ketone body, to acetoacetyl-CoA in the cytosol and plays an important role in lipid metabolism and in normal neuronal development in mice [7–9]. Our studies showed that ketone bodies not only are important alternative energy sources but also are crucial elements for lipid synthesis.

In energy production and lipid synthesis, acetyl units, such as acetyl-CoA and acetoacetyl-CoA, are important sources for various reactions and play an important role in a wide range of cellular processes. Many metabolic pathways, such as pyruvate catabolism, fatty acid oxidation, branched chain amino acid catabolism, ketolysis, ethanol/acetate metabolism, and citrate lysis generate acetyl-CoA [10]. Glucose is more easily oxidized than ketone bodies, however, a greater percentage of ketone bodies are utilized for cholesterol and fatty acids than for glucose [11]. Previous studies have shown that acetoacetate is a more efficient energy-yielding substrate for human mesenchymal stem cells than glucose [12]. These results suggest that metabolic substrate preference varies from cell to cell. Amino acids and fatty acids are used to produce acetyl CoA via glycogenesis and ketone body synthesis. Therefore, glucose and ketone bodies are the primary sources of acetyl units, which suggest that there is a correlation between glucose and ketone body metabolism. As such, Holleran et al. showed that when a high concentration of acetoacetate was provided to hepatoma cells, acetoacetate supplied 85% of the acetyl-CoA for lipogenesis, whereas only 2% was supplied by glucose [13]. Moreover, a recent study by Haydar et al. reported a link between insulin resistance and a single nucleotide polymorphism in the *AACS* gene (rs12818316) [14]. These results suggest that ketone bodies are key substrates for energy production and lipogenesis in hepatoma cells and AACS plays an important role in glucose metabolism.

A recent study showed that AACS is barely detectable in human liver [15]. However, some studies indicate that ketone bodies are utilized for lipid synthesis in cancer cells [16–18]. Moreover, we showed that the *AACS* gene is highly expressed in HepG2 cells (Fig. 1), suggesting that AACS plays a distinct role in hepatocarcinoma cells, especially during glucose metabolism, which is a common feature of cancer cells. While expression profile of human AACS mRNA in the normal tissues was well analyzed [19], a role of ketone body utilization by AACS in human and HepG2 remains unclear. In the present study, we determined whether a correlation between AACS and glucose metabolism exists in HepG2 hepatocellular carcinoma cells. We also examined whether the expression of AACS affects glucose uptake and ketone body metabolism in HepG2 cells. Finally, we investigated the effect of ketone bodies on glucose uptake and the effect of AACS knockdown on cell proliferation and sorafenib sensitivity.

**Figure 1.**
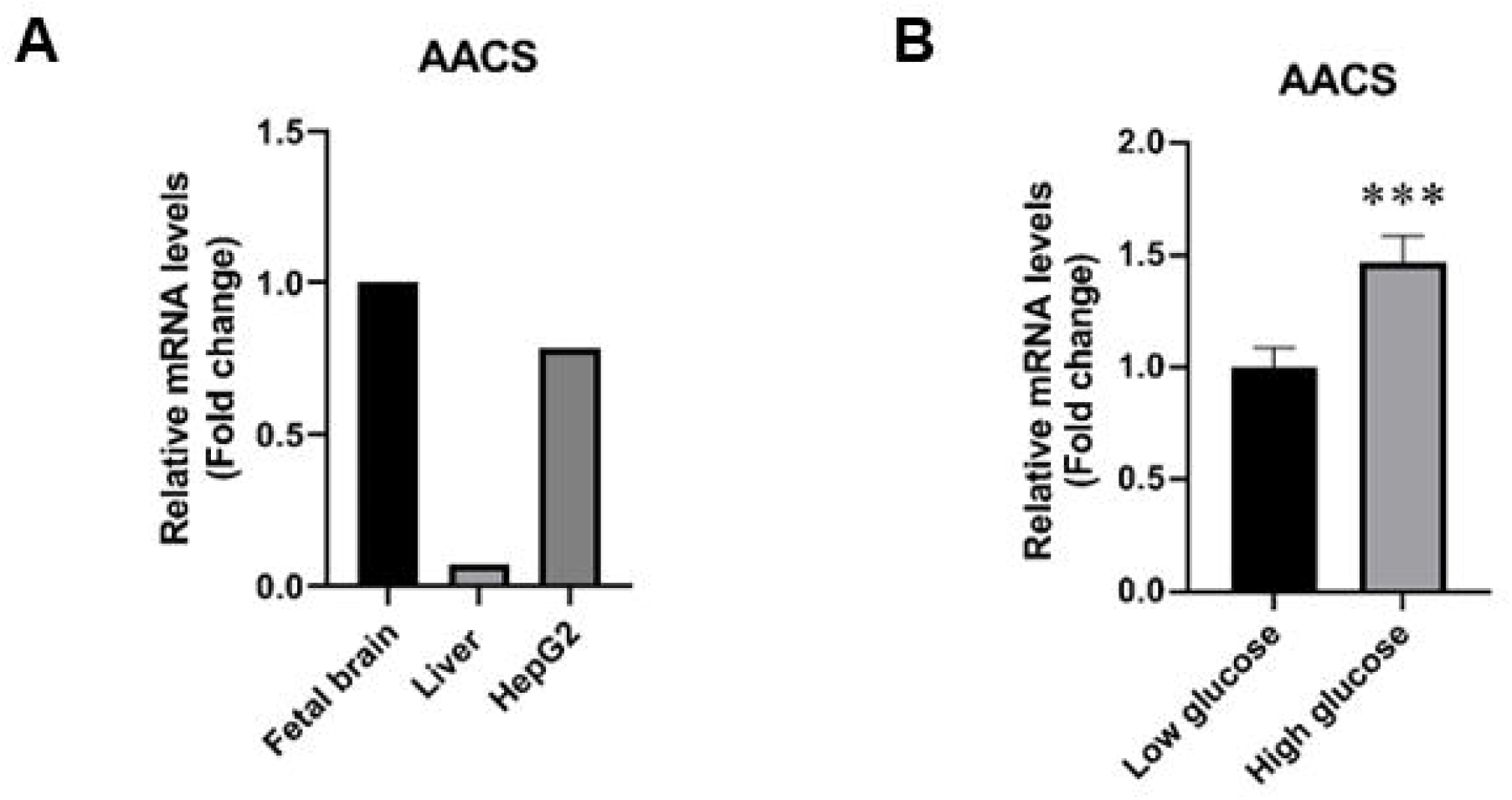
Effect of glucose concentration on the expression of AACS in HepG2 cells. (A) Expression pattern of AACS in human tissues and HepG2 cells. Total RNA from human fetal brain, liver, and HepG2 cells was reverse transcribed into cDNA. The mRNA levels of the AACS were measured by real-time PCR using specific primer sets. Gene expression was normalized to RPS18. The relative expression was quantified as foldchange compared with that of fetal brain. (B) HepG2 cells were incubated in either low glucose (1,000 mg/L) or high glucose (4,500 mg/L) medium. Total RNA was extracted and reverse transcribed into cDNA as described in the Materials and methods section. The mRNA levels of AACS and β-actin were measured using real-time PCR. Gene expression was normalized to β-actin expression. Relative expression was quantified as fold-change compared with low glucose treatment. Data are expressed as mean ± SD (n = 4). Comparing the high glucose group with the low glucose treatment group revealed the following: \*\*\**p*< 0.001.

## Results

### Effect of glucose concentration on the expression of AACS in HepG2 cells

A recent study showed that *AACS* is barely detectable in human liver [15]; however, some studies have indicated that ketone bodies are utilized for lipid synthesis in cancer cells [16–18]. Therefore, we investigated whether *AACS* is expressed in HepG2 human carcinoma cells. Consistent with our previous study, *AACS* was hardly detectable in human liver, but was highly expressed in HepG2 cells at a similar level as human fetal brain (Fig. 1A). This result suggests that AACS plays a distinct role in hepatocarcinoma cells, especially with respect to glucose metabolism, which is a common feature of cancer cells. We assessed *AACS* gene expression in the presence of low (1,000 mg/L) and high (4,500 mg/L) glucose concentrations to investigate whether glucose affects ketone body utilization (Fig. 1B). *AACS* mRNA levels were significantly elevated in the presence of high glucose concentration. These results suggest that ketone body utilization is regulated by glucose concentration in HepG2 cells and AACS plays an important role in glucose utilization.

### Effect of AACS knockdown on glucose uptake in HepG2 cells

Glucose uptake is the first step in glucose utilization, and tumor cells enhance glucose uptake and consumption for growth and proliferation. Therefore, elevated glucose uptake is a marker of tumors and an important target for cancer treatment [1]. We investigated whether AACS expression affects glucose uptake in HepG2 cells by infecting them with lentivirus encoding short hairpin (sh) RNA, targeting the *AACS* gene (shAACS). The AACS gene and protein expression were markedly decreased by treatment with shAACS (Fig. 2A and B). Moreover, glucose uptake in HepG2 cells was significantly reduced by AACS knockdown (Fig. 2C). These findings show that lipogenesis through ketone body utilization, which is regulated by glucose, affects glucose uptake.

**Figure 2.**
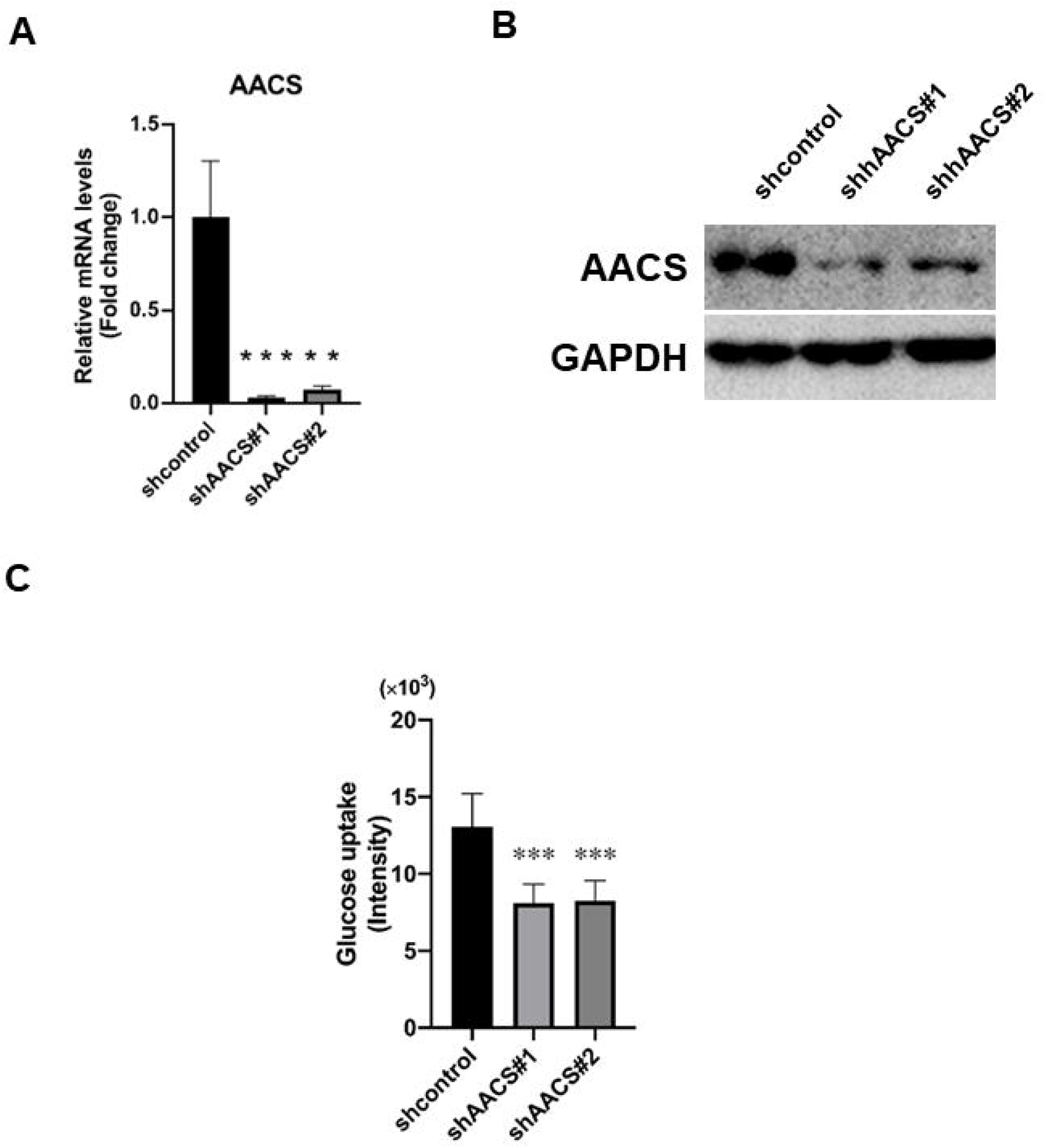
Effect of AACS knockdown on glucose uptake in HepG2 cells. (A) Effect of shRNA against AACS on the expression of AACS mRNA. HepG2 cells were infected with lentiviruses encoding shcontrol or shAACS. After 4 days, the culture media was replaced with high glucose DMEM, and the cells were incubated for 24 h. Then, the cells were harvested, and total RNA was extracted as detailed in the Materials and methods section. Total RNA was reverse transcribed into cDNA, and the mRNA levels of AACS and β-actin were measured by real-time PCR. Gene expression was normalized to β-actin expression. Relative expression was quantified as fold-change compared with the shcontrol group. Data are expressed as mean ± SD (n = 3). Comparing the shAACS groups with the shcontrol group revealed the following: \*\**p*< 0.01, \*\*\**p*< 0.001. (B) Effect of shRNA against AACS on the expression of AACS protein. HepG2 cells were infected with lentiviruses encoding shcontrol or shAACS. After 4 days, the culture media was replaced with high glucose DMEM. After 24 h, HepG2 cells were lysed in RIPA buffer. AACS and GAPDH protein expression was analyzed by western blotting with anti-AACS and anti-GAPDH antibodies, respectively. Lane 1: shcontrol, Lane 2: shhAACS#1, Lane 3: shhAACS#2. (C) Effects of AACS knockdown on glucose uptake in HepG2 cells. HepG2 cells were seeded, incubated for 24 h, and then infected with shAACS lentivirus. After 5 days, the cells were incubated with the fluorescent glucose analog, 2-NBDG (10 μM). The amount of 2-NBDG was measured by determining the fluorescence intensities at Ex/Em = 475/550 nm. Data are expressed as mean ± SD (n = 5). Comparing the shAACS groups with the shcontrol group revealed the following: \*\*\**p*< 0.001.

### Effect of AACS knockdown on the expression of lipogenic genes and glucose transporters

Next, we investigated whether AACS knockdown affects the expression of lipogenic genes and glucose transporters. We observed no difference in the expression of the lipogenic genes, *ACL, HMGCR*, or *FAS*, between shAACS-treated and shcontrol cells (Fig. 3A). Previous studies have shown that glucose uptake by GLUT1 and 9 is predominant in HepG2 cells [20]. The glucose transporters, GLUT1–4, and 9 were unchanged following treatment with shAACS (Fig. 3B). These results suggest that it is unlikely that genetic changes cause the observed changes in glucose uptake by shAACS treatment.

**Figure 3.**
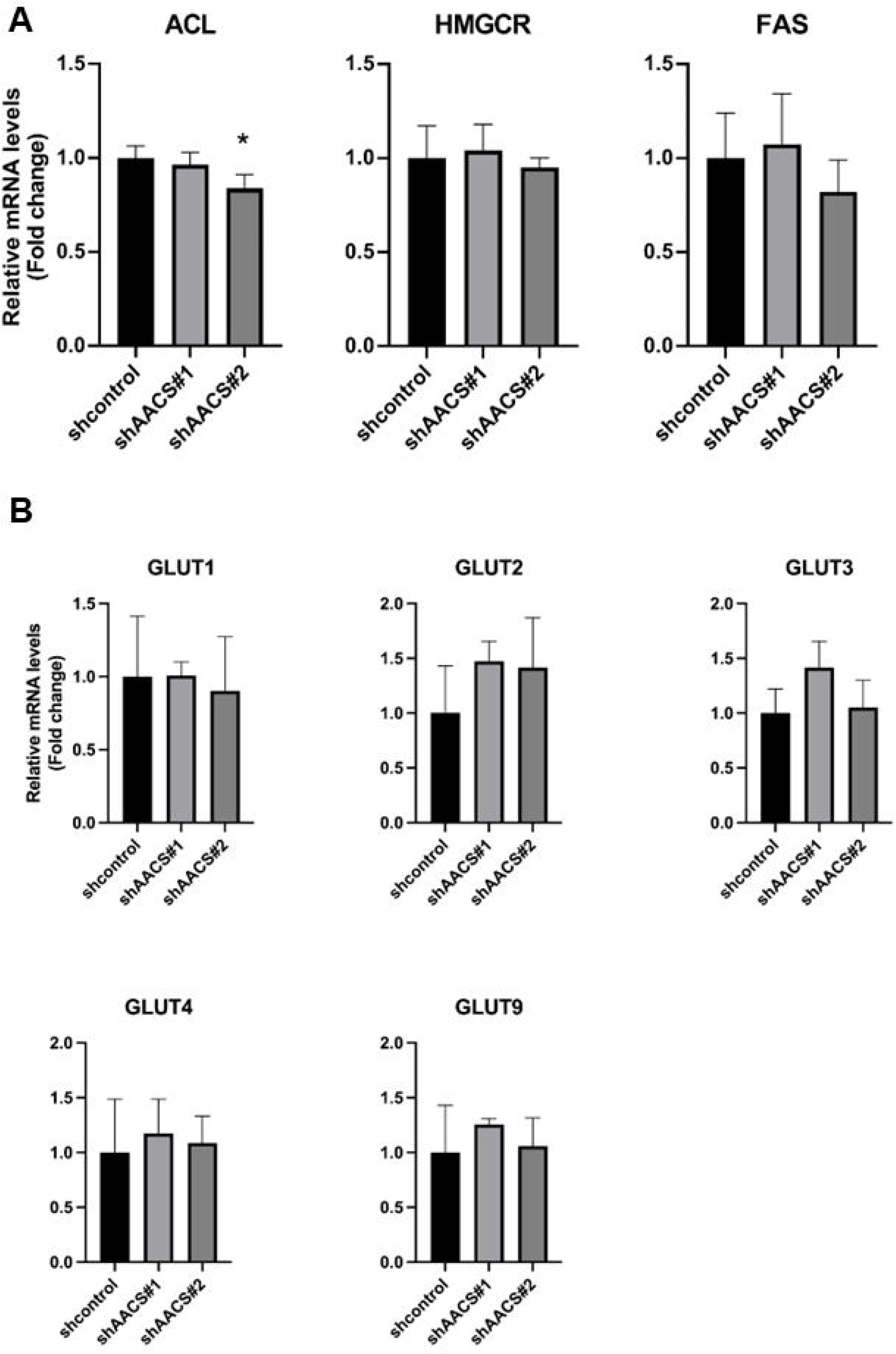
Effect of shRNA against AACS on lipogenic and transporter gene expression. (A) Effect of shRNA against AACS on lipogenic gene expression. HepG2 cells were infected with lentiviruses encoding shcontrol or shAACS. After 4 days, the culture media was replaced with high glucose DMEM. After 24 h, cells were harvested, and total RNA was extracted as detailed in the Materials and methods section. Total RNA was reverse transcribed into cDNA, and the mRNA levels of HMGCR, ACL, FAS, and β-actin were measured by real-time PCR. Gene expression was normalized to β-actin expression. Relative expression was quantified as fold-change compared with the shcontrol group. Data are expressed as mean ± SD (n = 3). Comparing the shAACS groups with the shcontrol group revealed the following: \**p*< 0.05. (B) Effect of shRNA against AACS on GLUTs. Total RNA was reverse transcribed to cDNA, and the mRNA levels of GLUT1–4, GLUT9, and β-actin were measured by real-time PCR.

### Effect of AACS knockdown on metabolite composition

Variation of metabolite composition in cells is an important factor in regulating glucose uptake [21,22]. AACS converts acetoacetate to acetoacetyl-CoA, which is incorporated into two essential metabolites, cholesterol and fatty acids. Therefore, we investigated whether knockdown of AACS affects the concentration of total cholesterol and NEFA in HepG2 cells. Cholesterol and NEFA were unchanged following treatment with shAACS (Fig. 4A). Ketone bodies represent an alternative energy source and are substrates for AACS. The effect of shAACS on the concentration of ketone bodies was examined (Fig. 4B). Knockdown of AACS increased total ketone body concentration in the media of HepG2 cells. Glucose and ketone bodies are important sources of acetyl units, suggesting that increased ketone body concentration may affect glucose uptake. Figure 4C shows that treatment of HepG2 cells with β-hydroxybutyrate significantly suppressed glucose uptake. These results suggest that the concentration of ketone body was increased by AACS knockdown, and that this metabolic change suppressed glucose uptake.

**Figure 4.**
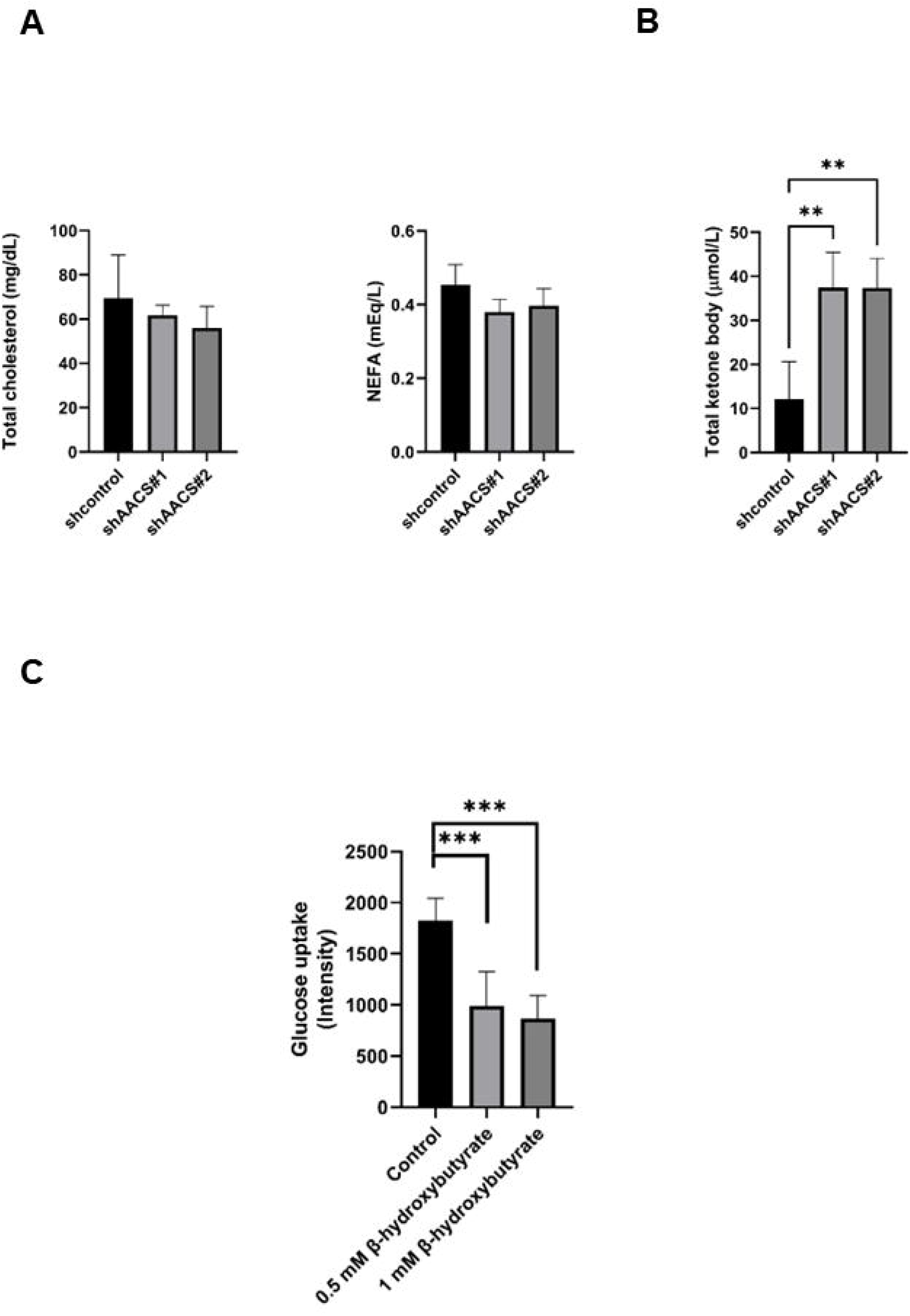
Effect of AACS knockdown on metabolite composition. (A) Effect of AACS knockdown on total cholesterol and NEFA. HepG2 cells were infected with lentiviruses encoding shcontrol and shAACS. After 4 days, the culture media was replaced with high glucose DMEM, and the cells were incubated for 24 h. The lipid fraction was then extracted using the Bligh and Dyer method. Total cholesterol and NEFA concentrations in HepG2 cells were determined using the LabAssay™ cholesterol and LabAssay™ NEFA. (B) Effect of AACS knockdown on ketone bodies in the media. The total ketone bodies in the media were assayed using the Autowako total ketone body assay kit. Comparing the shAACSs groups with the shcontrol group revealed the following: \*\**p*< 0.01. (C) Effect of β-hydroxybutyrate on glucose uptake. HepG2 cells were seeded into 96-well plates and incubated for 24 h. The cells were incubated with serum-free and phenol red-free DMEM for 3 h. Then, the cells were incubated with glucose-free DMEM for 15 min. Next, the cells were incubated with glucose-free DMEM containing the fluorescent glucose analog, 2-NBDG (10 μM) for 1 h. The amount of 2-NBDG fluorescence was measured. Comparing the shAACSs groups with the shcontrol group revealed the following: \*\*\**p*< 0.001.

### Effect of AACS knockdown on cell proliferation and sorafenib sensitivity

We investigated the effect of sorafenib on HepG2 cells and whether the knockdown of AACS affects cell proliferation (Figure 5). The knockdown of AACS led to lower prolifera-tive ability in untreated HepG2 cells. In sorafenib-treated cells, AACS knockdown was more effective in inhibiting cell proliferation than in untreated cells.

**Figure 5.**
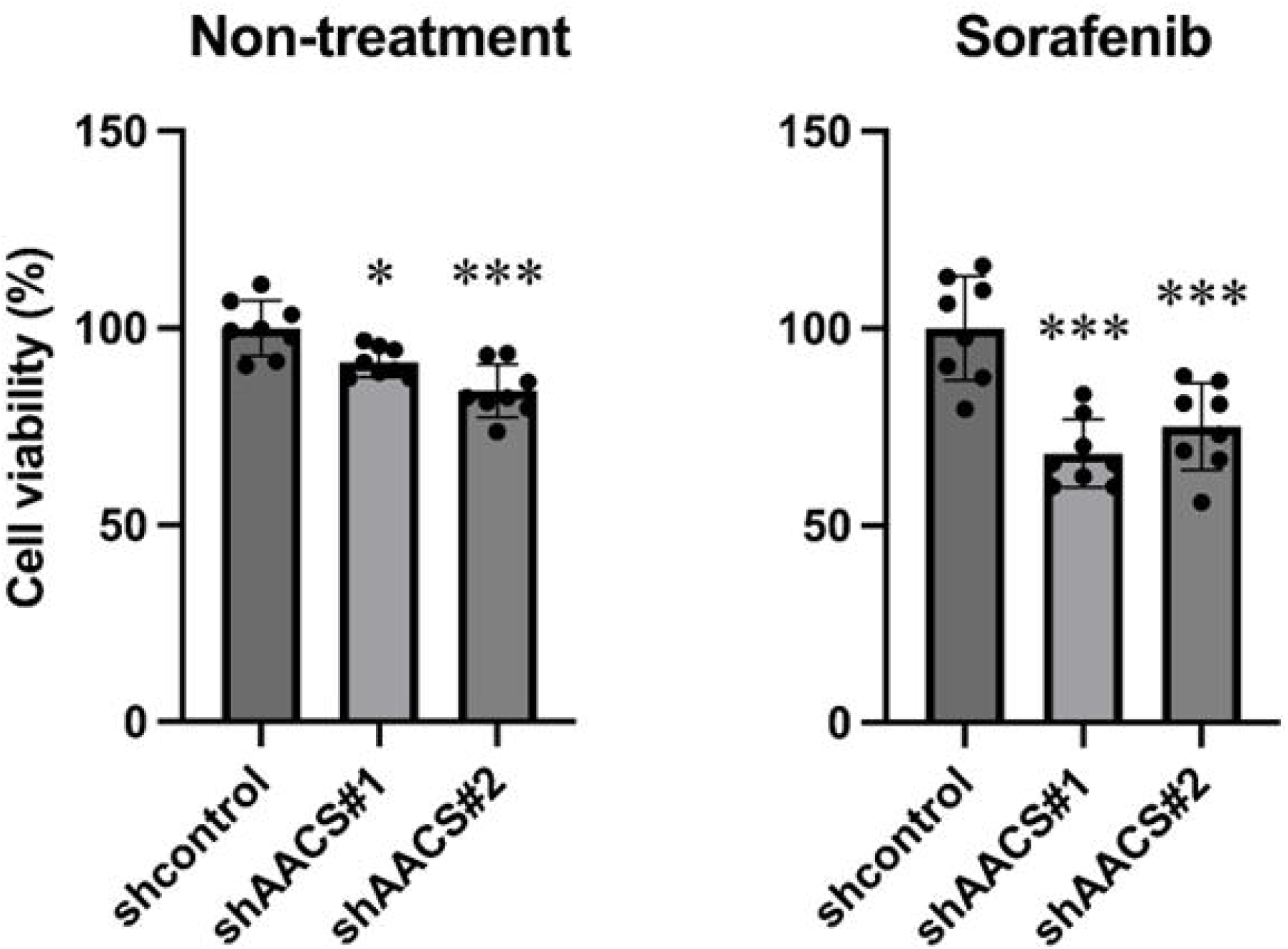
Effect of AACS knockdown on cell proliferation and sorafenib sensitivity HepG2 cells were infected with lentiviruses encoding shcontrol and shAACS. After four days, the cells were treated with 5 μM sorafenib for 24 h Cell viability of HepG2 cells was measured by MTT assay. Data are expressed as mean ± SD (n = 8). Comparing the shAACSs groups with the shcontrol group revealed the following: *p < 0.05, ***p < 0.001.

## Discussion

Previous studies and this study showed that *AACS* is barely detectable in human liver [15,19] but was highly expressed in HepG2 cells at a similar level as human fetal brain (Fig. 1A). We previously found that the liver contained high levels of AACS protein [7]. There is a difference in AACS expression in liver between human and mice. Previous studies showed that lipid metabolism in mice differs from that in humans [23–25]. Mice naturally lack expression of cholesteryl ester transfer protein (CETP), which shuttles lipids between lipoproteins. A previous study showed that the overexpression of simian CETP in mice protects against diet-induced insulin resistance [26]. Collectively, these results suggest that the expression of AACS may be related to differ-ences in lipid and glucose metabolism between human and mice, as well as between the liver of human and HepG2 cells.

Figure 1B shows that *AACS* gene expression was upregulated in the presence of high glucose concentration. A previous study identified 1,153 sites bound by carbohydrate response element binding protein and 783 target genes in the chromatin from HepG2 cells [27]. However, AACS was not included as part of the target genes, although AACS is upregulated by glucose as shown in Fig. 1B. Previous studies showed that the expression of mouse vesicular glutamate transporter 2 and the catabolic enzyme myo-inositol oxygenase are increased in the presence of glucose [28,29]. The interaction of Sp1 with their promoter regions increased in the presence of high glucose. These results indicate that Sp1 is an important mediator for glucose-stimulated transcriptional regulation. We and other authors previously reported that AACS is regulated by Sp1 in both mice and humans [30,31]. These results suggest that Sp1 is a key transcription factor for the upregulation of AACS in the presence of high concentrations of glucose.

We used NBDG to measure glucose uptake in HepG2 (Fig. 2C). NBDG is a fluorescent substance bound to glucose. Previous studies have shown that simultaneous treatment of the D-glucose inhibits the uptake of NBDG, but simultaneous treatment of the L-glucose does not inhibit uptake [20]. This result suggests that the behavior of NBDG is quite similar to that of glucose. As mentioned earlier, since GLUT1 expression is high in HepG2 cells, NBDG is thought to be preferentially taken up by GLUT1. Figure 2C shows that AACS knockdown decreased glucose uptake; however, the expression of glucose transporter genes was unchanged (Fig. 3B). These results suggest that some other factors affect glucose uptake. Therefore, we investigated the effect of AACS knockdown on metabolite composition, especially fatty acids and cholesterol. These lipids were unchanged (Fig. 4A) even though we previously showed that transient AACS knockdown in mouse liver resulted in decreased serum cholesterol [7]. The reasons for this may include species differences and metabolic changes in cancer cells. Knockdown of AACS increased the total ketone body concentration in the media of HepG2 cells (Fig. 4B). It is generally believed that ketone bodies are produced only when glucose is unavailable. However, these results suggest that ketone bodies are usually generated in HepG2, and that knockdown of AACS deviates ketone bodies from intracellular fluid.

Previous studies showed that ketonemia increased in glucose, glucose 6-phosphate, pyruvate, citrate, α-ketoglutarate, malate, and glutamate in the brain of rats. Acute ketonemia diminished flux through the hexokinase, phosphofructokinase, and pyruvate dehydrogenase reactions, but total influx of substrates into the Krebs cycle temporarily exceeded CO2 evolution, leading to increased concentrations of cycle intermediates [32]. Moreover, the ketogenic diet has a beneficial effect in treatment of GLUT1 deficiency [33]. These results suggest that the cells may be undergoing metabolic changes that do not require the use of glucose under high concentrations of ketone body. Also, a recent study has published that tyrosine kinase inhibitors affect the formation of the glucose transporter. Furthermore, we showed that acute exposure to ß-hydroxybutyrate reduced glucose uptake (Fig. 4C). This suggests that the effect of ketone bodies may be due to direct competition between glucose and ß-hydroxybutyrate as energy sources. The fact that simultaneous treatment of ketone bodies and glucose reduces glucose uptake suggests that ketone bodies may be more readily taken up than glucose via this mechanism [34].

Ketone bodies have a variety of roles in the cell, such as acting as signaling molecules. Knockdown of AACS increases intracellular ketone bodies, which leads to a shift in cellular energy metabolism toward ketone body consumption and decreased glucose uptake. The results also suggest that increased ketone bodies may increase concentrations of TCA cycle intermediates and increase intracellular energy metabolism. As cancer cells proliferate, the supply of energy disposition varies greatly depending on their location. Cancer cells located far from blood vessels are in a low glucose condition and need to generate their own energy, and ketone bodies may be useful for this purpose. In the future, a detailed study of energy metabolism in cancer cells will provide an important clue in further applications for cancer treatment and diagnosis.

Ketone bodies have beneficial effects such as tissue protection. A ketogenic diet, a high-fat, low-carbohydrate diet, supplies who has consumed it with ketone bodies, and they have been used for the treatment with [35]. Ketone bodies, generated through consumption of the ketogenic diet, have been identified as the main neuroprotective substance as treatment of neurons. β-Hydroxybutyrate and acetoacetate protected cells from oxidative stress [36,37]. Moreover, recent studies have shown that SGLT2 inhibitor upregulates ketone body production, resulting in protecting kidney from dysfunctions caused by diabetes [38]. Knockout of 3-hydroxymethyl glutaryl-CoA synthase 2 (HMGCS2), the rate-limiting enzyme for ketogenesis, attenuates renoprotective effect of SGLT2 inhibitor. These results suggest that ketone bodies have multi-dimensional roles in metabolism, tissue protection, and signal transduction [5].

Recent studies showed that ketone bodies weakened tumor cell viability and increased the survival rate of mice with systemic metastatic cancer [39]. Moreover, overexpression of HMGCS2 and enhanced ketone bodies production observed consequently help generate antiproliferative and antimigratory effects [40]. These results indicate that ketone body metabolism is an important target for cancer treatment. AACS was more highly expressed in cancer cells than in normal liver tissue (Fig. 1A) and was involved in ketone bodies’ deviation from cancer cells and glucose uptake in them (Fig. 2C and 4B). Furthermore, we found that knockdown of AACS, i.e., increased intracellular ketone body concentration, slightly attenuates cell proliferation and significantly increases sensitivity to the anti-cancer agent (Fig. 6). By studying this, we will be able to gain knowledge about the mechanism of cancer development and it’s treatment.

**Figure 6.**
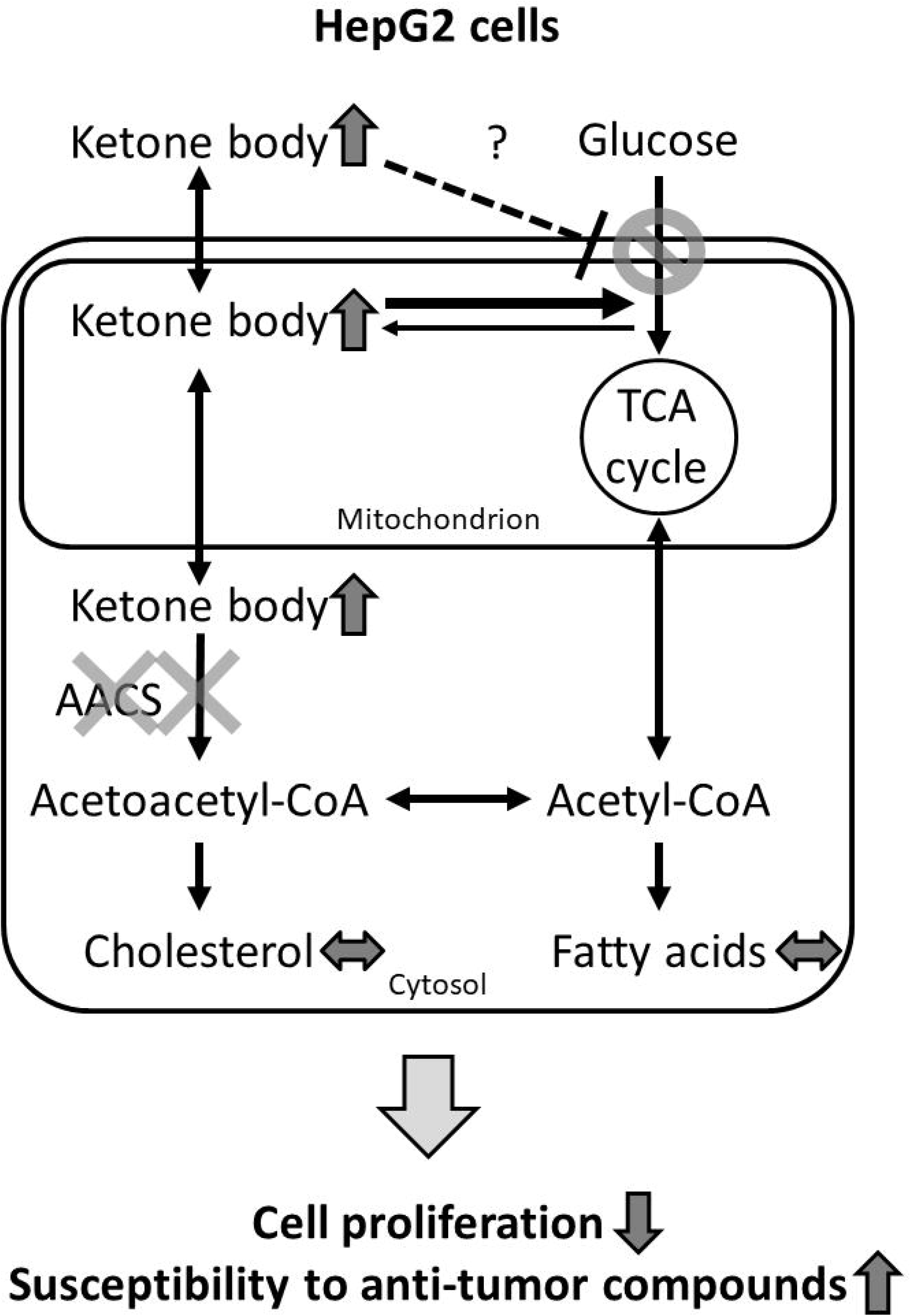
Proposed model for the effect of AACS knockdown on HepG2 cells

In the present study, we found that AACS mRNA levels are upregulated in the presence of high glucose concentration. AACS knockdown attenuated glucose uptake and increased the concentration of ketone bodies in the media. Moreover, treatment with β-hydroxybutyrate attenuated glucose uptake. Knockdown of AACS attenuated cell proliferation and significantly increased sensitivity to the anti-cancer agent. These results suggest that ketone body metabolism plays an important role in glucose uptake and AACS may be a viable candidate for understanding cancer metabolism and it’s treatment.

## Materials and Methods

### Real-time polymerase chain reaction

Total RNA from human liver and fetal brain was purchased from Clontech Laboratories, Inc. Total RNA (1 μg) was reverse transcribed with the PrimeScript™RT Reagent Kit (Takara Bio Inc.) according to the manufacturer’s protocol. For real-time polymerase chain reaction (PCR), the cDNA and gene specific primers were added to PowerUp SYBR™ Green Master Mix (Thermo Fisher Scientific) and subjected to PCR amplification in an Applied Biosystems StepOne system. The amplified transcripts were quantified using a standard curve with ribosomal protein S18 (RPS18) as an internal control. The real-time PCR primers were designed based on GenBank accession numbers using the Primer-BLAST tool available from the NCBI website [41]. The primer sequences were as follows: AACS: forward, 5’-ATGCTTGGCCGGAGTGAC-3’ and reverse, 5’-GATCACCCTCTCCTCCCTGT-3’ and ribosomal protein S18 (RPS18): forward, 5’-ATTAAGGGTGTGGGCCGAAG-3’ and reverse, 5’-TGGCTAGGACCTGGCTGTAT-3’.

### Cell culture

HepG2 hepatocarcinoma cells (JCRB1054) were maintained in Dulbecco’s Modified Eagle Medium (DMEM, 1,000 mg/L, low glucose) and supplemented with 10% fetal bovine serum (FBS; Gibco) at 37°C in a humidified atmosphere containing 5% CO_2_. For glucose stimulation, 2.5 × 10^5^ cells were seeded into a 6-cm dish. After 24 h, the culture medium was replaced with 10% FBS in DMEM (high glucose 4,500 mg/L or low glucose 1,000 mg/L), and the cells were incubated for 24 h. Total RNA was then extracted using a NucleoSpin^®^ RNA Plus kit (Takara Bio) and subjected to cDNA synthesis and real-time PCR as described above. The amplified transcripts were quantified using a standard curve with β-actin serving as an internal control. The primer sequences were as follows: β-actin: forward, 5’-GAGCACAGAGCCTCGCCTTT-3’ and reverse, 5’-TCATCATCCATGGTGAGCTGG-3’.

### Virus production and transduction

Virus production and transduction were performed as previously described [8]. Lenti-X 293 cells (Takara Bio Inc.) were seeded and transfected with pSIH-H1 shRNA lentivectors (System Biosciences) and packaging mix (Invitrogen) according to the manufacturer’s instructions. To facilitate AACS knockdown, HepG2 cells were infected with lentiviruses encoding shRNA sequences against two different human AACS coding regions as follows: shcontrol, CTTACGCTGAGTACTTCGA; shAACS #1, GGCGAGCTGGTGTGTACTA; and shAACS#2, CGGCAGCTCGGAAATCTAT. After 4 days, the culture media was replaced with high glucose DMEM, and the cells were incubated for 24 h. Total RNA was then extracted and subjected to cDNA synthesis and real-time PCR as described above. The primer sequences were as follows: ATP-citrate lyase (ACL): forward, 5’-GACTTCGGCAGAGGTAGAGC-3’ and reverse, 5’-TCAGGAGTGACCCGAGCATA-3’; HMGCR: forward, 5’-ATGGAAACTCATGAGCGTGGT-3’ and reverse, 5’-AGCTCCCATCACCAAGGAGT-3’; fatty acid synthase (FAS): forward, 5’-GACCGCTTCCGAGATTCCAT-3’ and reverse, 5’-GAGGCCTATCTGGATGGCAG-3’, glucose transporter (GLUT) 1: forward, 5’-GAACTCTTCAGCCAGGGTCC-3’ and reverse, 5’-ACCACACAGTTGCTCCACAT-3’, GLUT2: forward, 5’-CCCATCAATATTCAGCTGCCG-3’ and reverse, 5’-TGACAGTGAAAACCAGGGTCC-3’, GLUT3: forward, 5’-GACCCAGAGATGCTGTAATGGT-3’ and reverse, 5’-GACCCCAGTGTTGTAGCCAA-3’, and GLUT4: forward, 5’-CGACCAGCATCTTCGAGACA-3’ and reverse, 5’-CACCAACAACACCGAGACCA-3’, and GLUT9: forward, 5’-GTCACTGAGACCCATGGCAA-3’ and reverse, 5’-GCAG-GACCAGTCCTTTCTTCT-3’.

### Western blot analysis

Western blot analysis was performed as previously described [7]. Briefly, HepG2 cells were lysed in radioimmunoprecipitation buffer (RIPA) buffer [50 mM Tris–HCl (pH 8.0), 150 mM NaCl, 1 mM EDTA (pH 8.0), 1% Triton X-100, 1% sodium lauryl sulfate, and 0.1% sodium deoxycholate]. The lysed cells were centrifuged to remove debris (14,000*g* for 15 min at 4°C), and the resulting supernatants (cell lysate) were subjected to western blot analysis. Protein concentrations of the cell lysates were measured using a Bio-Rad protein assay kit [42], and 7.5 μg was separated using SDS polyacrylamide gel electrophoresis (7.5% gel). The separated protein was electrophoretically transferred to polyvinylidene difluoride membranes. The membranes were incubated with anti-human AACS [43] and GAPDH antibodies and then incubated with horseradish peroxidase (HRP)-conjugated secondary antibodies. HRP was detected with Immobilon Western Chemiluminescent HRP Substrate (Millipore).

### Measurement of glucose uptake

Glucose uptake was measured using 2-(N-(7-nitrobenz-2-oxa-1,3-diazol-4-yl)amino)-2-deoxyglucose (2-NBDG) [20]. HepG2 cells (2,000 cells) were seeded into 96-well plates and incubated for 24 h. The cells were then infected with shRNA against AACS (shAACS) lentivirus. After 4 days, the media was changed, and the cells were incubated for 24 h. The cells were incubated with serum-free DMEM. Then, the cells were washed with glucose-free DMEM once and incubated with glucose-free DMEM containing the fluorescent glucose analog, 2-NBDG (10 μM). After incubating for 24 h, the media was removed, and the cells were washed twice with 200 μL of Hank’s Balanced Salt Solution (HBSS). HBSS (200 μL) was added to the cells, and the amount of 2-NBDG was measured by determining the fluorescence intensities at Ex/Em = 475/550 nm using an ARVO X2 (PerkinElmer Co., Ltd.). To investigate the effect of β-hydroxybutyrate on glucose uptake, HepG2 cells (20,000 cells) were seeded into 96-well plates and incubated for 24 h. The cells were washed with serum-free and phenol red-free DMEM twice and incubated with serum-free and phenol red-free DMEM for 3 h. Then, the cells were washed with the glucose-free DMEM twice and incubated with glucose-free DMEM for 15 min. Next, the cells were incubated with glucose-free DMEM containing the fluorescent glucose analog, 2-NBDG (10 μM), with or without β-hydroxybutyrate. After incubating for 1 h, the media was removed, and the cells were washed twice with 200 μL of HBSS. Triton X-100 (0.1%) in H_2_O was added to the cells and subjected to three freeze–thaw cycles. The amount of 2-NBDG fluorescence was measured.

### Measurement of cholesterol, non-esterified fatty acids, and ketone bodies

HepG2 cells were infected with lentiviruses encoding shcontrol and shAACS. After 4 days, the culture media was replaced with high glucose DMEM, and the cells were incubated for 24 h. Then, the lipid fraction was extracted by the Bligh and Dyer method [44]. Total cholesterol and non-esterified fatty acids (NEFA) concentrations in HepG2 cells were determined using the LabAssay™ cholesterol and LabAssay™ NEFA (FUJIFILM Wako Chemicals) according to the manufacturer’s instructions. The total ketone body in the media was assayed using the Autowako total ketone body assay kit (FUJIFILM Wako Chemicals).

### Cell Viability assay

HepG2 cells (5000 cells/well) were seeded in 96-well plates and infected with lentiviruses encoding shcontrol and shAACS. After four days, the cells were treated with 5 μM sorafenib (Tokyo Chemical Industry Co., Ltd). After incubation for 24 h, the media were replaced with DMEM (1,000 mg/L, low glucose, and phenol-red free) supplemented with 10% FBS and 10% 5mg/mL 3-(4,5-dimethylthiazol-2-yl)-2,5-diphenyltetrazolium bromide (MTT) reagent (SIGMA) in PBS. After incubation for 3 h, the media were removed and 100 μL of dimethyl sulfoxide (Wako) was added to each well, and the absorbance at 570 nm was measured by a microplate reader.

### Statistical Analysis

All data are presented as the mean ± standard deviation (SD). The study results were analyzed using Graph Pad Prism software version 9 (GraphPad Software Inc.). Differences between individual groups were determined by repeated-measures two-way analysis of variance (ANOVA). A *p*-value <0.05 was considered statistically significant.

## Acknowledgments

This work was supported in part by a grant-in-aid from the Ministry of Education, Culture, Sports, Science and Technology of Japan (JSPS KAKENHI Grant Number 15K18911 and 18K14908).

## Notes

### Competing Interest Statement

The authors have declared no competing interest.

