## Supplementary figures and images for "Role of acetoacetyl-CoA synthetase in glucose uptake by HepG2 cells"

### Supplemental Figure 1

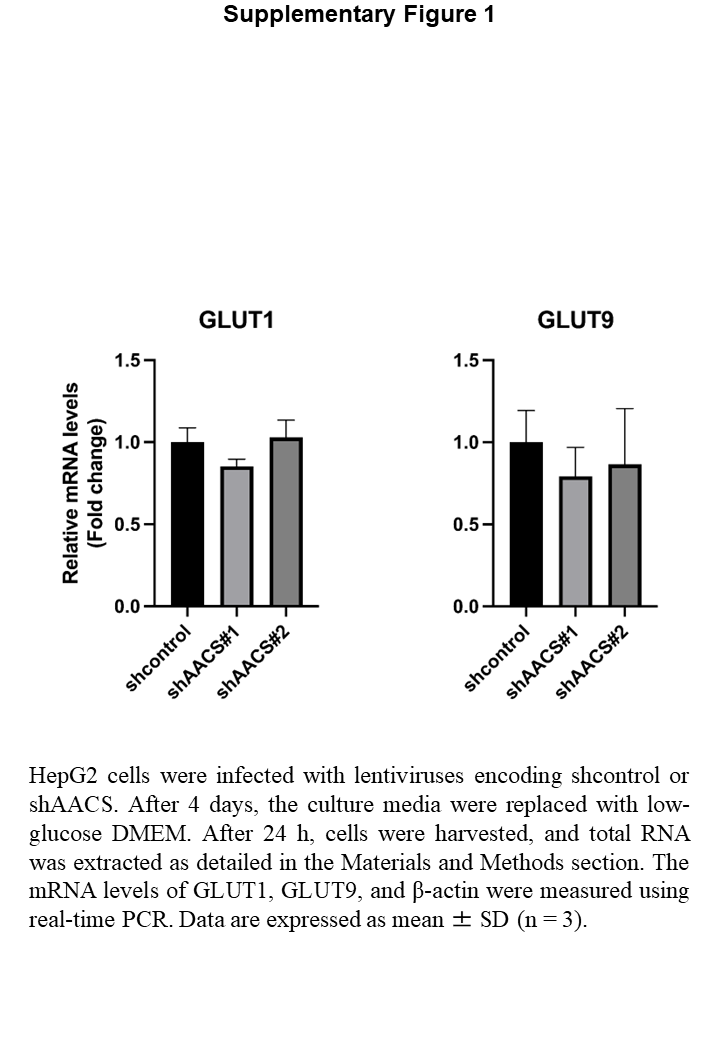
